# Preventing extinction in an age of species migration and planetary change

**DOI:** 10.1101/2023.10.17.562809

**Authors:** Erick J. Lundgren, Arian D. Wallach, Jens-Christian Svenning, Martin A. Schlaepfer, Astrid L.A. Andersson, Daniel Ramp

## Abstract

International and national conservation policies almost exclusively focus on conserving species in their historic native ranges, thus excluding species that have dispersed on their own accord or have been introduced by people. Given that many of these ‘migrant’ species are threatened in their native ranges, conservation goals that explicitly exclude these migrant populations may overlook opportunities to prevent extinctions and respond dynamically to rapidly changing environmental and climatic conditions. Focusing on terrestrial mammals, we quantified the extent to which migration, in this case via introductions, has provided new homes for threatened mammal species. We then devised alternative scenarios for the inclusion of migrant populations in mainstream conservation policy with the aim of preventing global species extinctions and used spatial prioritization algorithms to simulate how these scenarios could change global spatial conservation priorities. We found that 22% of all identified migrant mammals (70 species) are threatened in their native ranges, mirroring the 25% of all mammals that are threatened. Reassessing global threat statuses by combining native and migrant ranges reduced the threat status of 23 species (∼33% of threatened migrants). Thus, including migrant populations in threat assessments provides a more accurate assessment of actual global extinction risk among species. Spatial prioritization simulations showed that reimagining the role of migrant populations to prevent global species extinction could increase the importance of overlooked landscapes, particularly in central Australia. Our results indicate that these various and non-exhaustive ways to consider migrant populations, with due consideration for potential conservation conflicts with resident taxa, may provide unprecedented opportunities to prevent species extinctions. We present these alternatives and spatial simulations to stimulate discussion on how conservation ought to respond, both pragmatically and ethically, to rapid environmental change in order to best prevent extinctions.

## Introduction

The redistribution of organisms through human introductions has provided opportunities for a number of species outside their historic ranges. Many of these species are threatened or extinct in their native ranges while thriving in their introduced ones, presenting a conservation paradox (*1–3*). This process of biotic redistribution is expected to accelerate, both from continuing globalization, but also as species migrate in response to ongoing landscape and climatic changes. However, migrant species, both those introduced by humans and many of those who dispersed on their own (e.g., *4*, *5*), are widely considered pests and thus excluded from biodiversity datasets and threat assessments (*6*). The result is that they are almost universally targeted by conservation eradication and control programs, regardless of whether they are endangered or extinct in their native ranges (*7*).

Preventing extinction is one of conservation’s primary aims (*8*). However, this aim can come into conflict with mainstream preferences for preserving only historically native life (*9*). These conflicts raise important questions about whether conservation biology should respond to widespread ecological and climatic changes by broadening its valuation of organisms beyond those restricted to their historically native ranges. Take the Javan rusa deer (*Rusa timorensis*) as an example. The Javan rusa is threatened by poaching and habitat loss in its native range of Indonesia (*10*). However, the Javan rusa has established a successful population in continental Australia after introduction by humans in the late 1800s (*11*). Conservation’s primary response to the Javan rusa, and all other deer in Australia (including other threatened ones), is to eradicate them (*12*, *13*). However, what happens if the Javan rusa becomes extinct in their native range while the Australian population lives on? Under existing protocols, would the International Union for the Conservation of Nature (*10*) list the Javan rusa as ‘extinct in the wild’ or ‘extinct’ while Australia continued with its eradication plans? A progressive alternative would be to engage in exploratory discussions of how conservation policies could attach value to introduced Javan rusa, either by accommodating their presence in their new range as part of biotic reorganization under planetary change, or as a source population for future repatriation to Java.

For the most part, conservation has set aside these paradoxes (but see *14*) due to claims that introduced organisms have fundamentally different—and unwanted—effects relative to native ones (*15*, *16*). While we acknowledge that in specific cases (especially on islands) introduced organisms have contributed to extinctions, meta-analyses have repeatedly shown that it is impossible to distinguish between native and introduced organisms on the basis of their effects (*17–21*); that some ‘invasive’ organisms targeted for eradication turn out to be native endemic species (*22*); that introduced organisms are not a leading cause of biodiversity loss (*23*); and that when we look we find that many introduced organisms sustain ecosystem services and facilitate other species (*24–28*). While there remains disagreement about these points among conservation scientists, we cannot ignore the fact that continued exclusion and eradication of introduced populations may exacerbate extinction of vulnerable groups of plants and animals at a time of monumental planetary change.

The Earth is undergoing a radical pace of landscape transformation, primarily from agriculture and development (*29*), with significant future deforestation projected to be driven by increasing meat consumption rates (*30*). Likewise, some climate warming projections set us on a course to the Early Eocene (*31*), when Northern Greenland was a warm-temperate forest (*32*, *33*). While the concept of a native range is already fraught by ambiguities (*4*), this concern only escalates when organisms are no longer able to live in their historic distributions. For conservation policy to respond proactively, compassionately, and pragmatically to these changes, we must anticipate ways to value biodiversity in a time of species redistribution, both for species introduced by humans and those migrating on their own accord (*34*). Accounting for migrant species in conservation is not to dismiss potential conflicts with resident taxa but allows us to see the ecological processes and conservation opportunities flourishing in the increasingly novel environments of our time.

To provide inspiration for these discussions we quantified various ways the biotic redistribution of threatened species could be accounted for, focusing on mammals as an example taxonomic group. Given the scarcity of data on species that have migrated on their own (but which are often considered non-native or ‘invasive’ as well, e.g., rusty crayfish (*Orconectes rusticus*) and Cattle Egret (*Bubulcus ibis*) in North America, *4*, *5*), we focused on wild populations of ‘migrants’ whose ranges have expanded through human introductions. We then proposed three alternative scenarios of how we might value migrant biodiversity with the aim of preventing global extinctions: *conservative*–regarding migrant populations as refuges and applying existing native-range-based threatened status across the full range; *expansive*–reassessing threat status based on the full range (migrant plus native) and thus reducing the threat status of species globally; and *independent*–acknowledging the non-redundant value of native populations and the emerging evolutionary trajectories of migrant populations, thus assigning threat status to native and migrant populations separately. To understand how these differing approaches could influence conservation policy we tested the relative effect of these various formulations using quantitative spatial prioritization simulations to evaluate future global conservation goals that may be implemented to reduce extinction risk.

## Methods

We focused on terrestrial mammals (*n*=1,225 species) as their threat statuses are well known and their native distributions have been thoroughly mapped by the IUCN Red List (2018, *10, 35*). These ranges were added to introduced migrant ranges digitized from the peer-reviewed literature, government reports, newspaper articles, and a variety of databases (Table S1). To describe the overall pattern of modern biotic redistribution we quantified those biogeographic realms (*36*) that have donated versus received migrant mammals and what percentage of threatened species per mammalian family and order have migrant populations.

### Alternative scenarios

We formulated three scenarios based on different formulations of how conservation might include migrant biodiversity, *conservative*, *expansive*, and *independent*, and compared these to the current *native-only* approach. In the *native-only* scenario–the *status quo*–we assigned conservation value only to populations of threatened species within their native ranges (Near Threatened species were considered threatened) based on the normative premise that non-native populations have little, no, or negative value. However, following IUCN Red List guidelines (section 2.1.3, *10*), migrant populations introduced with the intent of reducing extinction risk (e.g., conservation translocations) were treated as native populations, as were migrations geographically adjacent to native populations (*10*).

In the *conservative* scenario, the Red List threat status is established solely on the basis of the native populations. The threat status is then applied to the entire species’s range (the area of the combined native and migrant ranges) even if the migrant population was larger than the native population. This was done under the normative premise that introduced populations could one day acquire a value-status equivalent to native populations, but also acknowledges that redistributed populations have not stood “the test of time” in their new regions and might therefore be unstable and collapse (*37*), and that global stochasticity in human pressures may warrant the most conservative approach to prevent global extinctions.

In the *expansive* scenario, we reevaluated species threat statuses globally, based on the mammals full current range (the area of the combined native and migrant ranges). By doing so, some species would be down-listed or delisted globally, deprioritizing both the native and migrant ranges. This scenario is based on the normative premise that migrant populations are considered legitimate components of biodiversity and are thus monitored and protected with equal care across their range. To reassess threat statuses, we used IUCN listing criteria and assumed a linear relationship between range size and population size (following *38*). Utilizing the IUCN Red List listing criteria, we considered a 20% change in total range size relative to their native-only range size as criteria for one step-change in threat level (e.g., from Critically Endangered to Endangered).

Finally, in the *independent* scenario we assigned conservation value to migrant populations of species independent of native populations on the normative premise that both native and introduced populations have their own evolutionary trajectories and thus unique and equal value. Under the *independent* scenario, valuing migrant populations did not affect the threat status of native populations (unlike in the *expansive* scenario). The threat status of migrant populations was based on the total migrant range size relative to the native range size (**Table 1**).

**Table 1.**
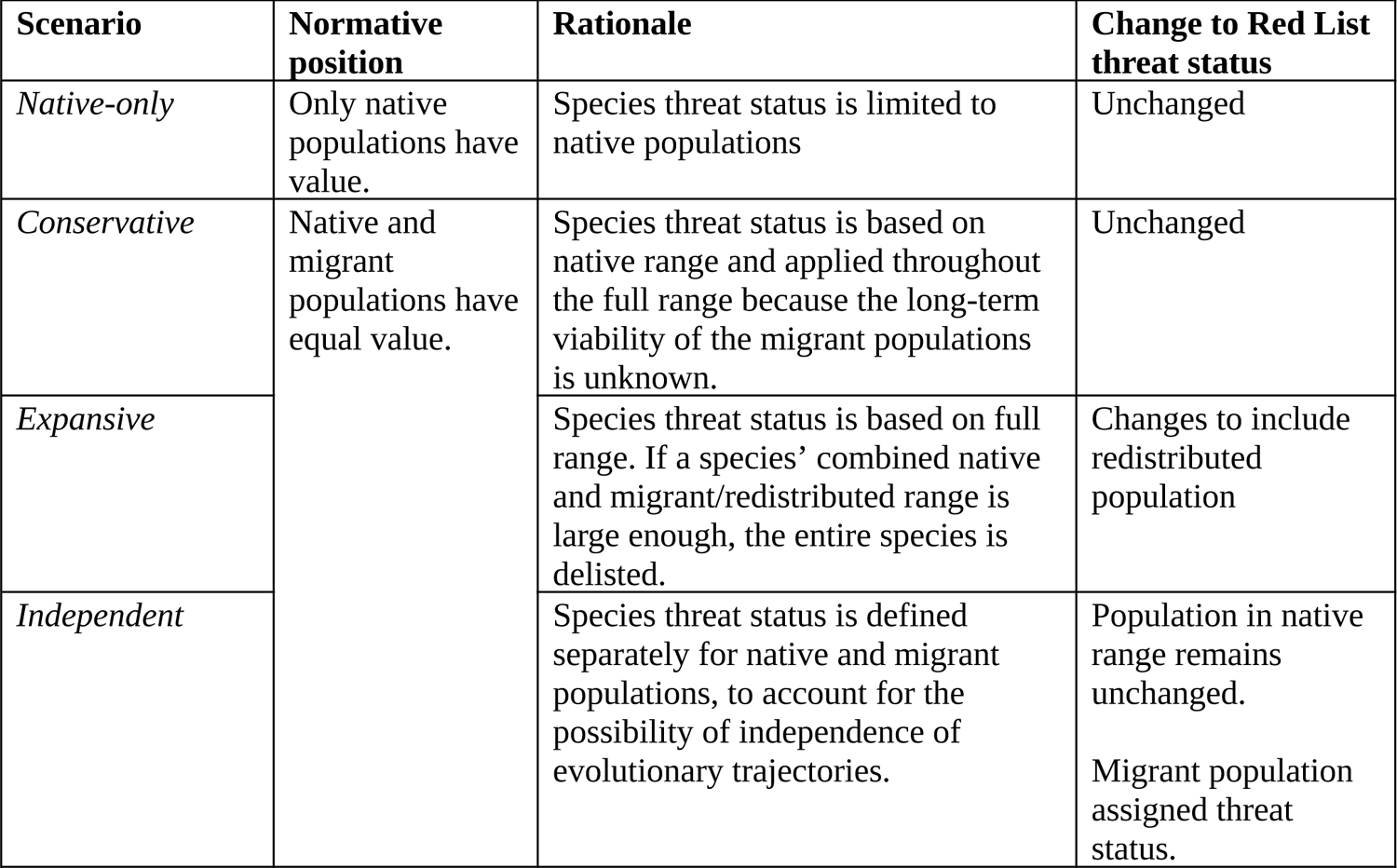
Value scenarios. Alternative ways to imagine the exclusion or inclusion of migrant organisms under the umbrella of conservation concern. Table lists the scenarios analyzed, their rationale, and changes (if any) to threat statuses. Changes in threat status (in expansive and independent scenarios) affected simulations both by removing species (if delisted) and by changing priority weighting (see below).

| **Scenario** | **Normative position** | **Rationale** | **Change to Red List threat status** |
| --- | --- | --- | --- |
| *Native-only* | Only native populations have value. | Species threat status is limited to native populations | Unchanged |
| *Conservative* | Native and migrant populations have equal value. | Species threat status is based on native range and applied throughout the full range because the long-term viability of the migrant populations is unknown. | Unchanged |
| *Expansive* |  | Species threat status is based on full range. If a species' combined native and migrant/redistributed range is large enough, the entire species is delisted. | Changes to include redistributed population |
| *Independent* |  | Species threat status is defined separately for native and migrant populations, to account for the possibility of independence of evolutionary trajectories. | Population in native range remains unchanged.<br>Migrant population assigned threat status. |

### Spatial prioritization

To gauge how these various scenarios might alter potential conservation action we conducted spatial prioritization algorithms to identify areas of high conservation importance globally for each scenario. To do so, we rasterized species ranges to produce feature layers for prioritization analysis using the R package ‘*exactextractr’* v0.4.0 (*39*) with a Mollweide projection at a 30×30km resolution. Migrant ranges reported at the country scale or within provincial boundaries (n=12 populations of 8 species) were omitted from spatial prioritization analyses because the large sizes of these political entities would lead to global delisting, even if populations were low. However, to account for these populations (many in central and east Asia), 1% of that total area was used in re-assessing species threat statuses for the *expansive* value scenario. This cutoff is arbitrary yet conservative for the purposes of this simulation.

We conducted spatial prioritization analyses using the R package ‘*prioritizr’* v5.0.1 (*40*), which uses integer linear programming techniques to find optimal solutions for spatial conservation planning problems. We used a ‘maximum utility’ objective to find the ‘biggest bang for the buck’ solution that most efficiently conserved as many species as possible per a specified conservation budget, in this case the number of land units (e.g., pixels).

Species were assigned weights based on their threat status. We weighted Near Threatened species with a weight of 1, Vulnerable species with a weight of 3, Endangered species with a weight of 5, and Critically Endangered, Extinct in the Wild, and Extinct species with a weight of 7. Thus, the prioritization algorithm gave extra importance to protecting the most endangered taxa. In doing so, changes in threat status in the *expansive* and *independent* value scenarios altered the importance of those populations in each prioritization simulation. We iteratively calculated prioritization solutions for each value scenario, increasing the total number of land units in the conservation budget from 1% of the Earth’s surface to 30%. The resulting solutions were summed to provide a continuous ranking of relative priority per land unit.

## Results

### Biotic redistribution

We identified 265 mammal species with at least one migrant (introduced) population. Of these, 70 (22%) are threatened in their native ranges, mirroring the 25% of all terrestrial mammal species that are threatened (*10*). Migrants that have been introduced into different realms originate from all realms bar Antarctica and Oceania and have most commonly been donated from the Indomalaya (59 species, 32.1% of redistributed mammals) and Palearctic (46, 25.0%), followed by the Afrotropics (31, 16.8%), the Nearctic (26, 14.1%), Australasia (11, 6.0%), and the Neotropics (11, 6.0%) (**Fig. 1A**). These species were received primarily by the Palearctic (57, 18.0%), Australasia (54 species, 17.0%), the Neotropics (51, 16.1%), the Nearctic (47, 14.8%), followed by Afrotropics (39, 12.3%), Oceania (32, 10.1%), Indomalaya (29, 9.1%) and Antarctica (8, 2.5%) (**Fig. 1A**).

**Figure 1.**
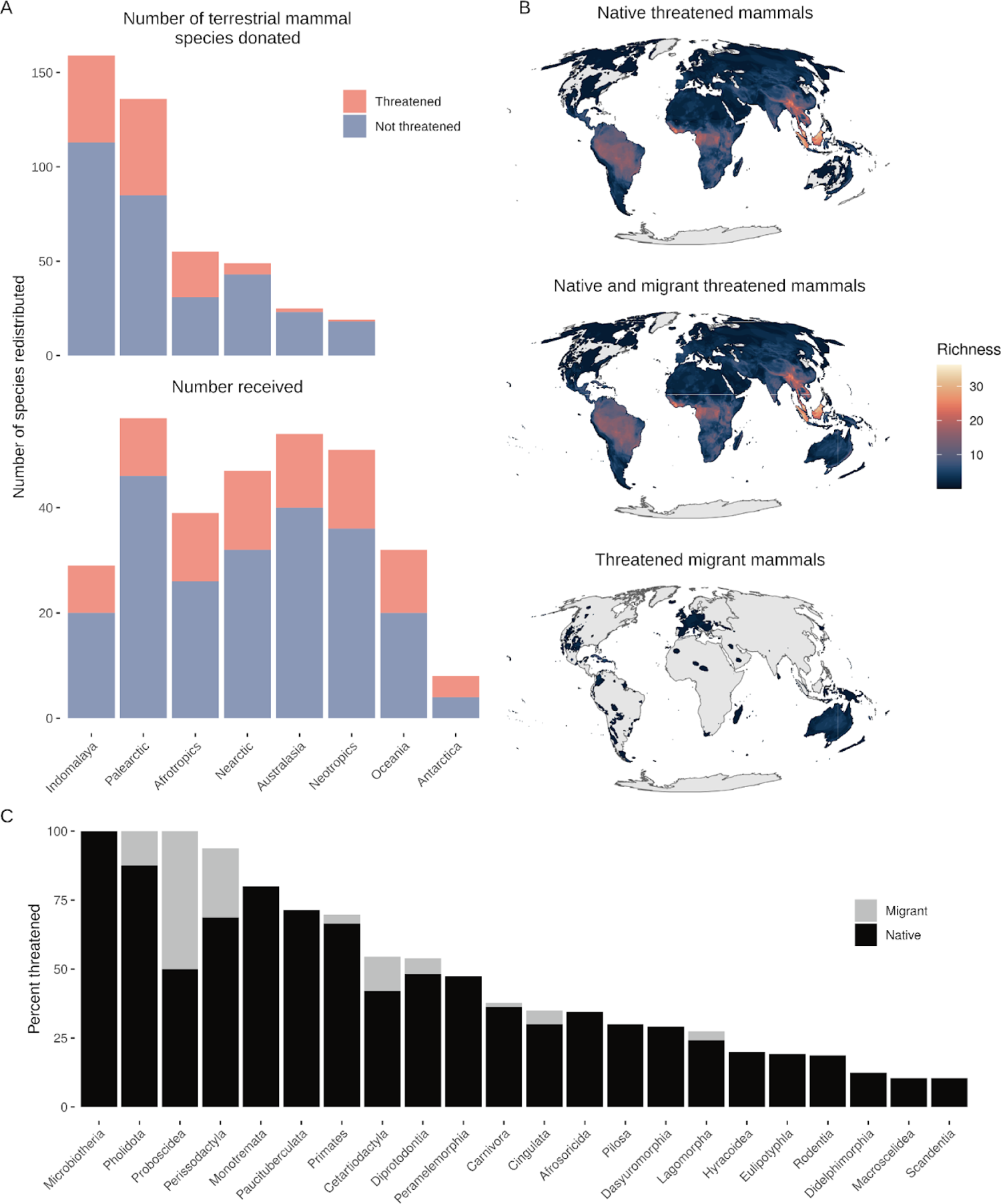
Overview of modern biotic redistribution of terrestrial mammals. **A**. Number of migrant mammals donated by biogeographic realms (top) versus received by realms (bottom). Fill indicates whether the migrant is considered threatened or not in their historic native range. **B.** Species richness of threatened mammals when only considered native mammals (IUCN Red List) and when considering all threatened mammals, including migrants. Migrant population distributions themselves are shown in the lowest map panel. **C.** Percent of mammal orders listed as threatened with fill indicating species that have migrant populations (see Fig. S2 for percent of mammal families).

Migrants that are threatened in their native range were donated mostly from Indomalaya (16 species, 32.6%) followed by the Palearctic (14, 28.6%), Afrotropics (13, 26.5%), Neartic (3, 6.1%), Australasia (2, 4.1%), and Neotropics (1, 2.0%) (**Fig. 1A**). These species were received by the Neotropics (15 species, 16.1%), Neartic (15, 16.1%), Australasia (14, 15.1%), and Afrotropics (13, 14.0%) followed by Oceania (12, 12.9%), Palearctic (11, 11.8%), Indomalaya (9, 9.7%) and Antarctica (4, 4.3%).

Biotic redistribution has slightly increased threatened mammal species richness in Australia, southwestern Nearctic, the Caribbean, and the Nearctic (**Fig. 1B**). Migrant mammals represent threatened species from 9 of 22 terrestrial mammal orders (**Fig 1C**) and 52 of 115 families, including up to 100% of threatened species in some families (**Fig S1**).

### Conservative scenario

Only valuing native populations and excluding redistributed migrant ones (*native-only scenario*) establishes tropical parts of South America, Africa and Asias as the top priority for most effectively protecting the maximum number of threatened mammals per land area (Fig. 2A). However, if migrant populations of threatened mammals are ascribed equivalent conservation value (*conservative scenario*), Australia—home to 16 threatened migrant mammals—becomes almost equally important for global conservation of threatened mammals (Fig. 2A-B). Likewise, the Caribbean and parts of South America increase in priority while parts of Africa and central Asia become slightly less emphasized (**Fig 2B**).

**Figure 2.**
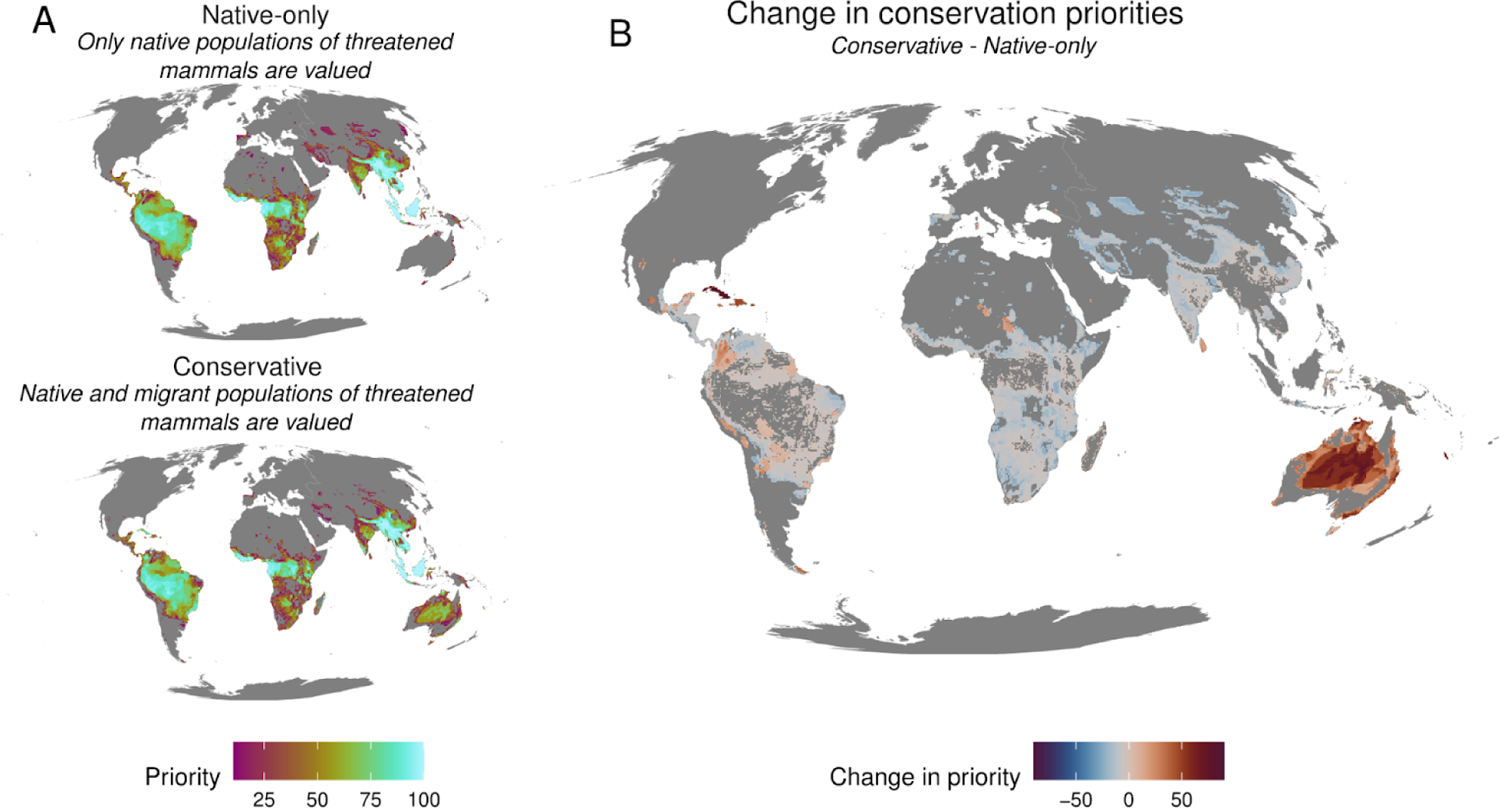
Including migrant populations of threatened species leads to an increased prioritization of Australia and the Caribbean, both of which possess a diversity of migrant species that are threatened in their native ranges.

### Expansive scenario

Of the 70 threatened migrant mammal species, redistribution has extended their total range by an average of 781% (between 0.01% and 18,000%). Reassessing the global threat status of these mammals based on their entire range (e.g, the *expansive* value-scenario) reduces the threat status of 23 (∼33%) of these threatened mammal species, of which 20 (∼29%) would become least concern (Fig. 3A).

**Figure 3.**
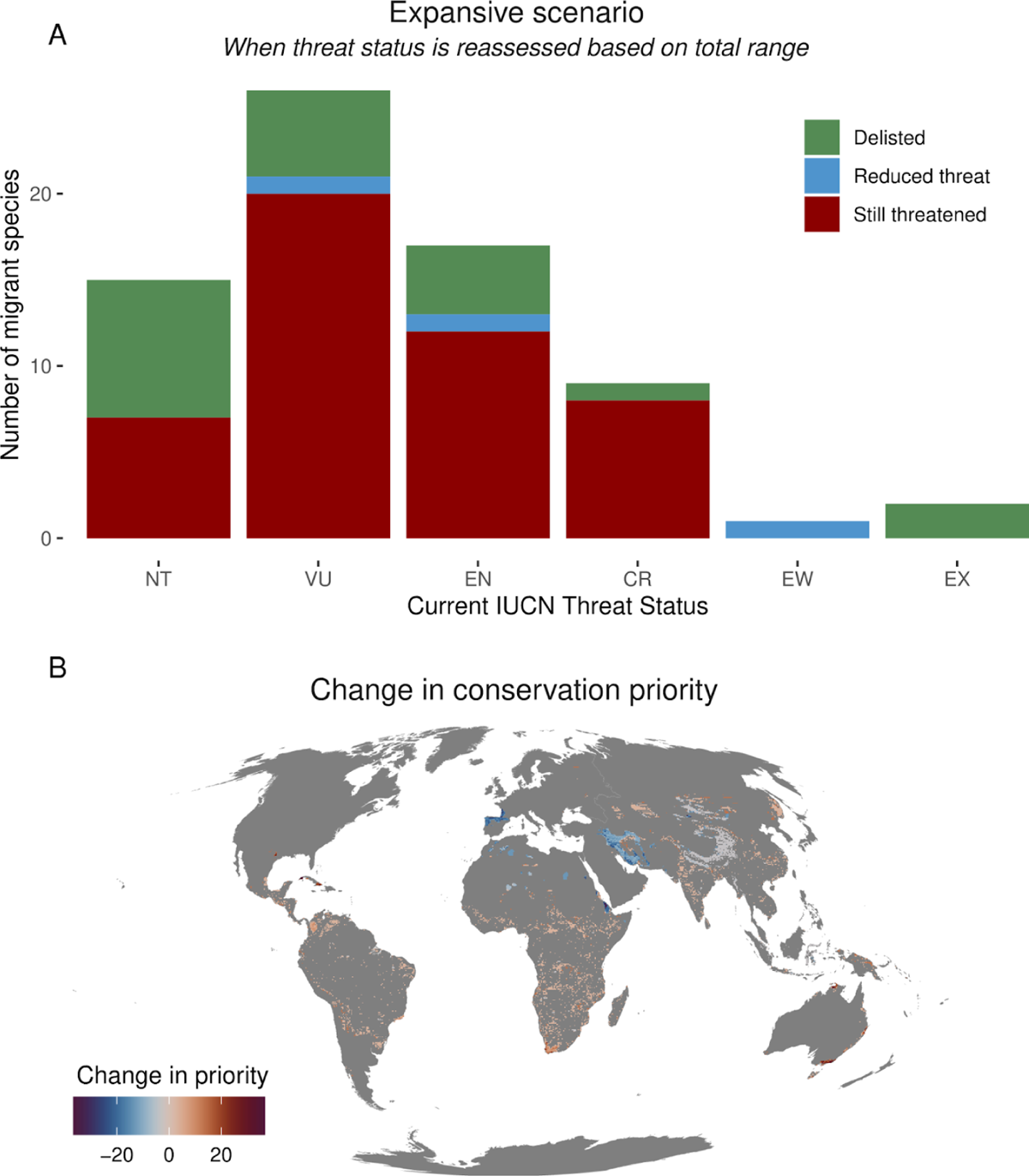
Assessing global threat status based on combined native and migrant (i.e., introduced) ranges delists 20 threatened mammal species and reduces threat statuses for 3 others (**A**). Including migrant populations in conservation policy thus reveals important differences among species in their exposure to global extinction. (**B**) Doing so leads to limited shifts in conservation priorities towards those species for whom there is no safety net.

In this scenario, spatial prioritization solutions show little difference from *native-only* solutions, except for the deprioritization of limited areas in Europe and central Asia due to the delisting or reduction in threat of these species. This leads to increased prioritization among taxa that do not have migrant populations; thus, conservation resources would flow to those species that have not found refuge through global redistribution (Fig. 3B). Given the constraints of finite conservation resources, this scenario provides a clear policy pathway for preventing further mammal extinction by enabling the redirection of resources to those who need it most.

### Independent scenario

Finally, a third scenario might retain the threat status of native populations of species but consider the status of migrant populations separately, based on an independent assessment of threat and population resilience. This scenario thus argues that modern migrant mammals are valuable, both as refuges for species threatened in their native ranges and in their own right, while maintaining that native populations are valuable in a non-redundant way to migrant ones. Importantly, this scenario reframes the narrative of conservation from one dedicated to recreating historic configurations to a futurist forward-looking pursuit where conservation works to promote adaptation and flourishing while preventing extinction under planetary change.

Under this scenario, 12 of the 70 migrant species had large enough distributions to be treated as Least Concern and could thus be omitted from spatial prioritization (17% of threatened migrant mammals), while 58 migrant species were considered threatened based on their small migrant range size, thus increasing the total number of threatened and (and thus prioritized) mammal species by 2.7% (Fig. 4A). Prioritizing the conservation of these migrant populations alongside native populations increased the conservation importance of parts of the Nearctic (Texas and the Caribbean), the Neotropics, southeastern Australia, and Europe (Fig. 4B).

**Figure 4.**
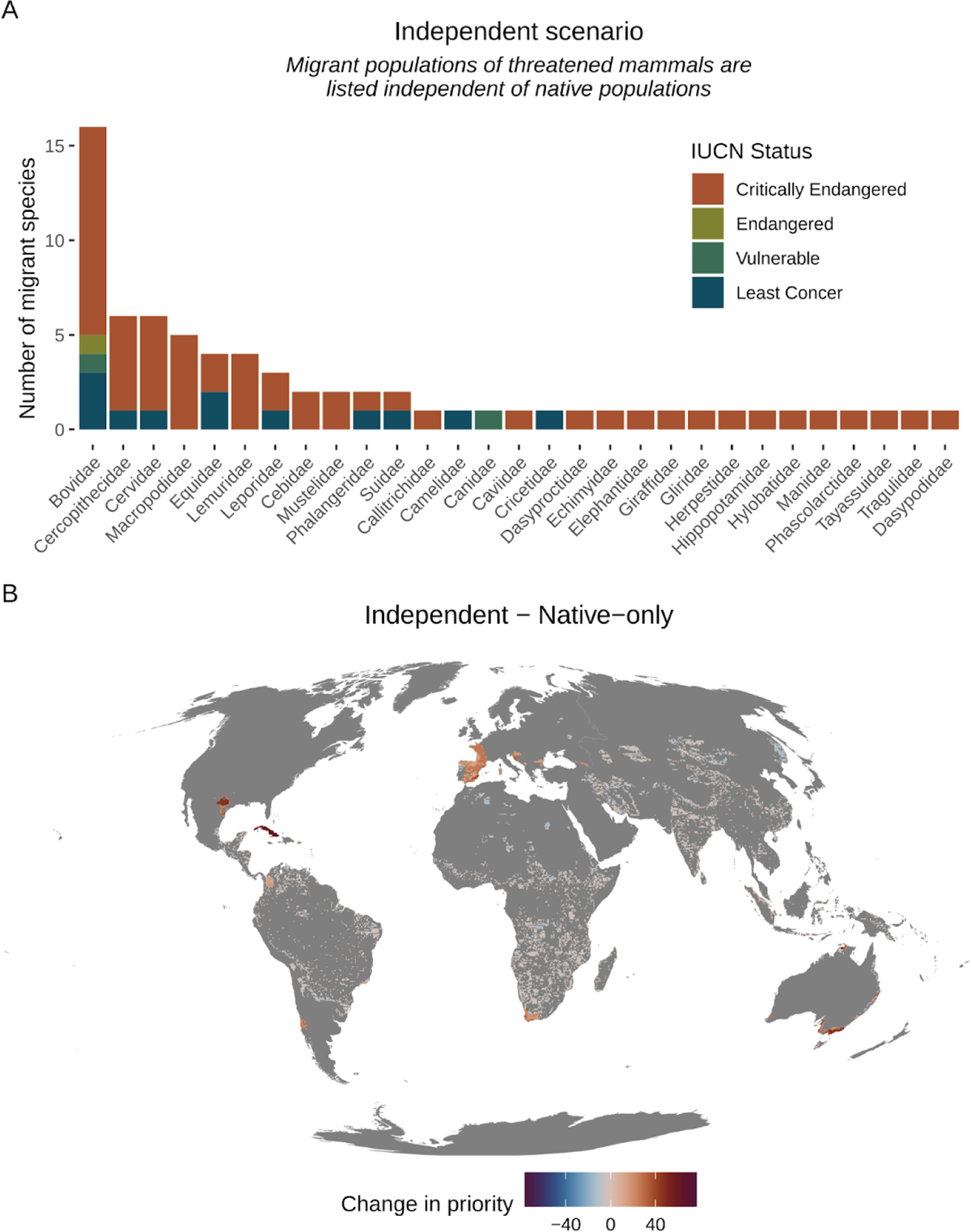
Prioritizing distinct populations of migrant species for their own evolutionary and intrinsic value adds new distinct, and threatened, populations per mammal family (**A**) but has little effect on spatial conservation priorities (B).

Overall, we found that various methods to ascribe value to migrant threatened mammal species led to shifts in how we might best prioritize conservation efforts to prevent extinctions. Relisting mammals to include migrant ranges (either changing global threat status or adding migrant populations as their own valued entities) led to only mild changes in the total number of threatened mammals overall (Fig. 5A), reflective of the broad scale of global mammal endangerment (*41*). However, including migrant populations did have consequences for the percentage of species’ ranges that were prioritized (Fig. 5B), especially for species currently listed as Extinct or Extinct in the Wild. Doing so shifted conservation priorities spatially, increasing the importance of landscapes in Australasia and the Nearctic (Fig. 5C). Importantly, protecting threatened species in their migrant ranges, which are often in wealthier regions, may help share some of the burden of species conservation with those who can best afford it (*42*).

**Figure 5.**
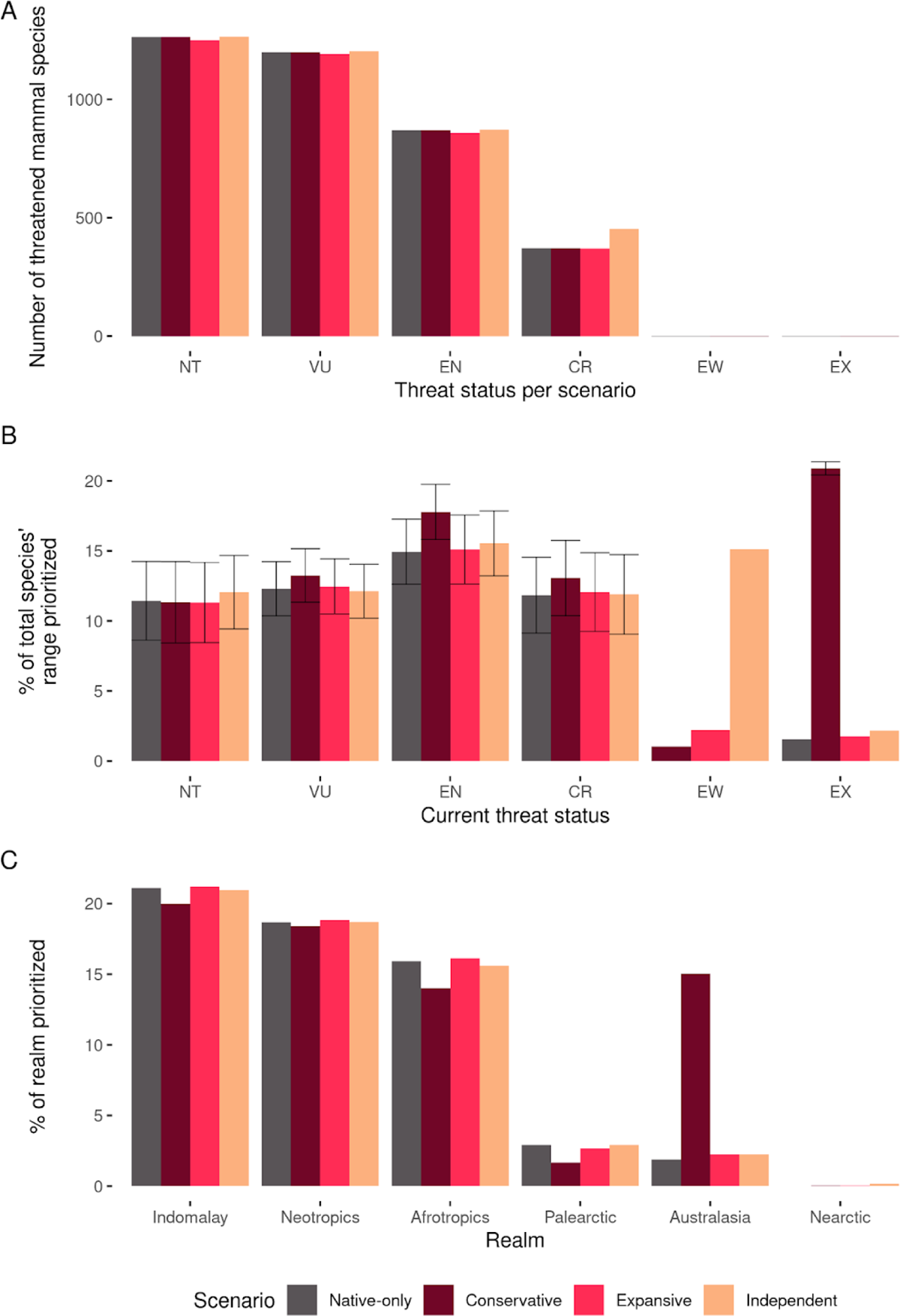
The influence of including migrant populations in global prioritization scenarios. **A**. Re-assessing mammal threat status based on the combined migrant and native population sizes (*expansive* scenario) or independently listing migrant populations (*independent* scenario) led to only modest shifts in the number of mammals threatened. **B.** Including migrant populations changes the percent of threatened species’ ranges that are prioritized, especially for species currently considered extinct in the wild or extinct. **C.** Including migrant populations shifts conservation priorities towards wealthier biogeographic realms, which may help shoulder the cost of species conservation towards those who can afford it best (*42*).

## Discussion

In an age of extinction and ecosystem change, we explored how different conservation policy avenues towards migrant populations could change conservation goals. While the redistribution of biota is fundamental to evolutionary diversification and potentially to ecosystem resilience (*43*), human-assisted migration of species is generally considered aberrant and harmful. The consequence is that most national and international conservation policies, including frameworks like the IUCN Red List, ignore migrant biodiversity (*6*), rendering populations of 70 of the world’s threatened mammals invisible (*44*). While reconsidering the conservation status of migrant populations of threatened species does not radically alter the global conservation outlook for most terrestrial mammals, which have not found refuge through introduction, some important opportunities for discussion are revealed.

For example, we found that if migrant populations were considered as important as native populations, then the rich migrant large herbivore (megafauna) community of central Australia could be considered a major conservation priority, whose protection could reduce the risk of extinction of eight of these ecologically important species and their globally endangered functional group (*1*, *45–48*). Among these migrant populations of Australian megafauna are the world’s only population of wild dromedary camel (*Camelus dromedarius*), which went extinct in the wild in its native range 3,000-5,000 years ago and is therefore not listed on the IUCN Red List, as well as feral water buffalo (*Bubalus arnee bubalis,* critically endangered in their native range) and feral donkeys (*Equus asinus africanus,* critically endangered, Fig. 6A). Many of these populations retain genetic diversity lost from their native conspecific populations (*14*, *49*, *50*) and thus, genetically, may be a lifeline to the persistence of these species globally. Moreover, active conservation efforts in the native ranges of some of these species is difficult, if not impossible; for example, the remaining 23-200 African wild asses occur in areas currently occupied by warring groups (*10*).

**Figure 5.**
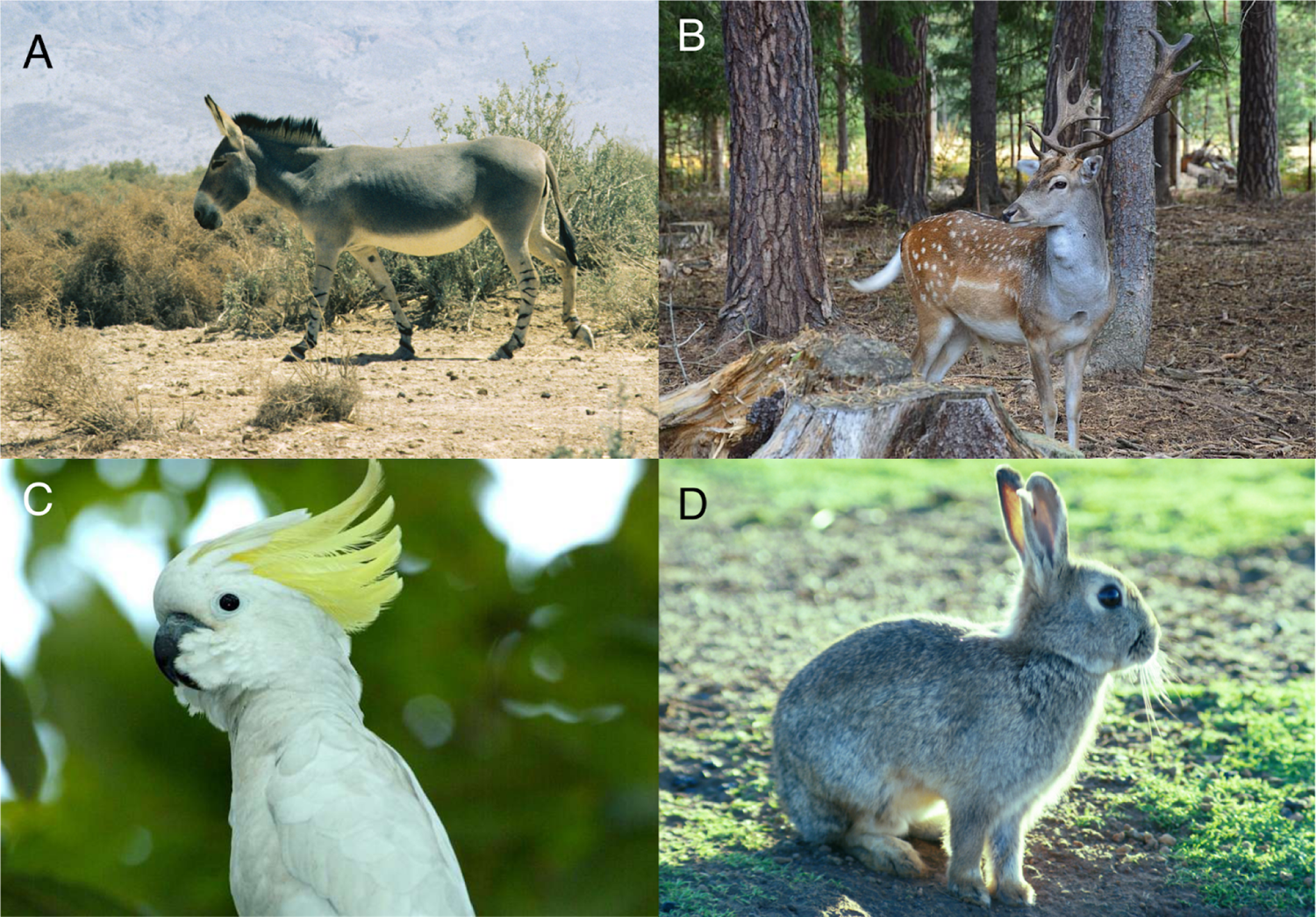
Threatened species whose conservation may benefit from inclusivity towards migrant populations. **A.** The African wild ass (*Equus africanus*) is Critically Endangered with a population of 23-200 individuals in remote, war-torn areas of Northeast Africa. Yet their migrant post-domestic relatives have populations in North and South America and Australia. **B.** Only 400-500 fallow deer (*Dama dama*) live in their native range yet they are ranked Least Concern to the IUCN because Roman-era redistributions across western Europe and Great Britain have been culturally adopted as native. **C.** There are at least 187 recorded migrant populations of European rabbits (*Oryctolagus cuniculus*), which are Endangered in their native range. **D.** The yellow-crested cockatoo (*Cacatua sulphurea*) is critically endangered in Indonesia, where its populations have collapsed from the pet trade (*58*). Yet, in one pet-trade hub, Hong Kong, they now have a thriving and genetically-diverse population (*59*).

Despite prevailing assumptions about the impacts of these species, to the best of our knowledge, the only published empirical research on their ecological effects in Australia has shown that while these animals do cause trampling and reduce cover of preferred foods (as with any megafauna) (*51*, *52*), they also reduce wildfire frequency and intensity, which increases tree survivorship and growth rates (*45*, *53*), and that they maintain desert surface water availability which in turn sustains endemic fish populations (*54*). Given their unique influences on ecosystem functioning and their global endangerment, megafauna are key targets for rewilding efforts around the world, which includes the intentional introduction of feral megafauna (*55*). Yet for the most part, migrant mammals have been excluded from this vision (*56*). In an age of change, perhaps the safest route to ensure a future with megafauna, and their influences (*57*), would be to include these migrant populations by considering both their intrinsic value and their contribution to conservation agendas.

Valuing migrant populations by including them in global threat assessments (e.g., the *expansive* scenario) immediately delists 20 species and reduces threat levels for 3 others. While this is only 1.9% of all threatened mammals, the inclusion of migrant populations in global threat assessments reduces threat statuses for more species than the estimated 7-16 mammal extinctions prevented by active conservation interventions between 1993 and 2020 (*60*).

Including migrant populations in global threat assessments is not unheard of, particularly if the introduction is old enough and has become culturally adopted as native. For example, despite having a native population of <200 individuals, the fallow deer (*Dama dama*) is listed as Least Concern by the IUCN (**Fig 6B**) because of ancient introductions across western Europe (*10*). While this appears to contradict IUCN Red List guides for the inclusion of introduced populations, as these introductions lacked ‘conservation intent’ and the migrant range exceeds ‘reasonable’ geographic proximity (guidelines section 2.1.3, IUCN Red List 2018), this establishes a useful precedent for engaging in discussions of how migrant populations can be considered to re-examine IUCN listings.

The third scenario we examined considered the threat statuses of migrant populations independently from native populations (e.g., the *independent* scenario). This scenario maintained conservation concern for native populations while protecting the emerging eco-evolutionary trajectories of migrant ones (*61*). Rapid evolution within migrant populations—and interacting native ones—can conceive new taxa and ecological interdependencies (*62–68*). Dispersal, including large-scale interchanges from the geologic collisions of continental and oceanic biotas, is the progenitor of biodiversity and potentially of ecological resilience (*43*). Instead of describing these processes as ‘unnatural’, this scenario values the emergence of new ecologies and evolutionary trajectories. For example, the European rabbit (*Oryctolagus cuniculus*) (**Fig 6C**) is classified as endangered (*10*) but has established at least 187 distinct populations, ranging from the subarctic to the tropics and across all 16 of the world’s biomes (as defined by *36*). Charles Darwin initially considered one island population of European rabbits to be a new species due to their remarkably divergent morphology (*69*). Should consideration be made for the future biodiversity of *Oryctolagus*, particularly given the threat status of European rabbits in their native range?

These are not straightforward debates, as the example of the European rabbit highlights. Australian farmers have long struggled to reduce rabbits, investing considerable resources to deliver technological solutions aimed at their eradication (*70*). In the radically altered landscapes of the Anthropocene, emerging ecologies may be unpredictable and may increase conflict. While this may be true in some circumstances, what is clear is that current policy frameworks require reimagining to promote futures that incentivize coexistence, improve equity and justice for all species, and prevent further extinction and biodiversity loss. Ultimately, the exclusion of migrant populations does not serve conservation universally well and establishes narratives that mirror modern-day politics that demonize the redistribution of people (*71*). The challenge for conservation is to rise above historical valuations of belonging to find ways to promote safety for non-human life wherever it finds itself (*34*).

This may also include a broadening of conservation concern to include overlooked, modified, and urbanized landscapes—conservation frontiers outside traditional wilderness models where many threatened migrant populations reside. In addition to mammals, numerous threatened birds have established migrant populations, especially in urban environments (Fig. 6D). In many cases, the very same process endangering their native populations—the wildlife trade—is the source of these new populations (*72*). Expanding conservation efforts into these anthropogenic environments provides novel opportunities to protect species without land acquisition, to find common ground with environmental justice efforts in urban areas, and to connect the populace, for whom nativeness may not be a shared value, with caring for the organisms with whom their lives intersect (*73*).

Global ecosystems, climate, and society are dynamically shifting, giving rise to often unpredictable and unprecedented challenges. Excluding modern migrant biodiversity from conservation policy may inhibit our ability to respond creatively to these and other changes. We have presented three alternative scenarios as a means to investigate how valuing migrant species in different ways might influence global conservation priorities and frameworks like the IUCN Red List. We do not necessarily endorse any of these potential alternatives to native-only conservation. Indeed, it may be possible to conceive of how policy frameworks might utilize input from all these scenarios (and others we have not considered) to arrive at context-dependent decision making that best protects ecosystems and prevents further extinctions. Together, our results suggest that stewarding migrant populations of threatened species, with due consideration for potential conservation conflicts with resident taxa, may make important contributions to preventing global extinctions.

## Acknowledgments

We thank Martin Jung for advice regarding spatial prioritization methods and Mark Davis for feedback on earlier drafts of this manuscript. EJL was funded by the International Postgraduate Research Scholarship from the University of Technology Sydney. ADW considers this work a contribution to her Australian Research Council Future Fellowship. JCS considers this work a contribution to the Center for Ecological Dynamics in a Novel Biosphere (ECONOVO), funded by Danish National Research Foundation (grant DNRF173) and his VILLUM Investigator project “Biodiversity Dynamics in a Changing World”, funded by VILLUM FONDEN (grant 16549).

---

**Table S1.**
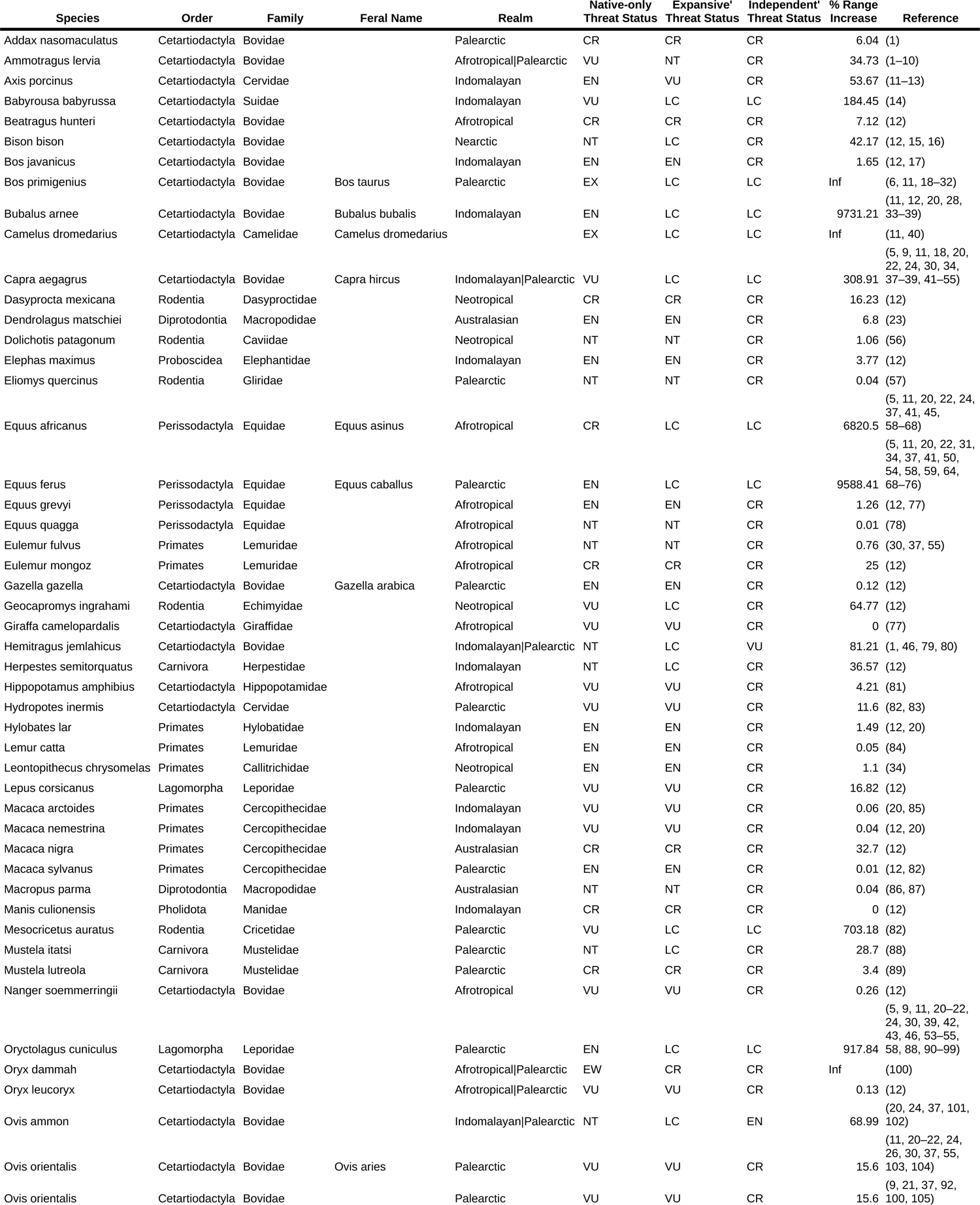
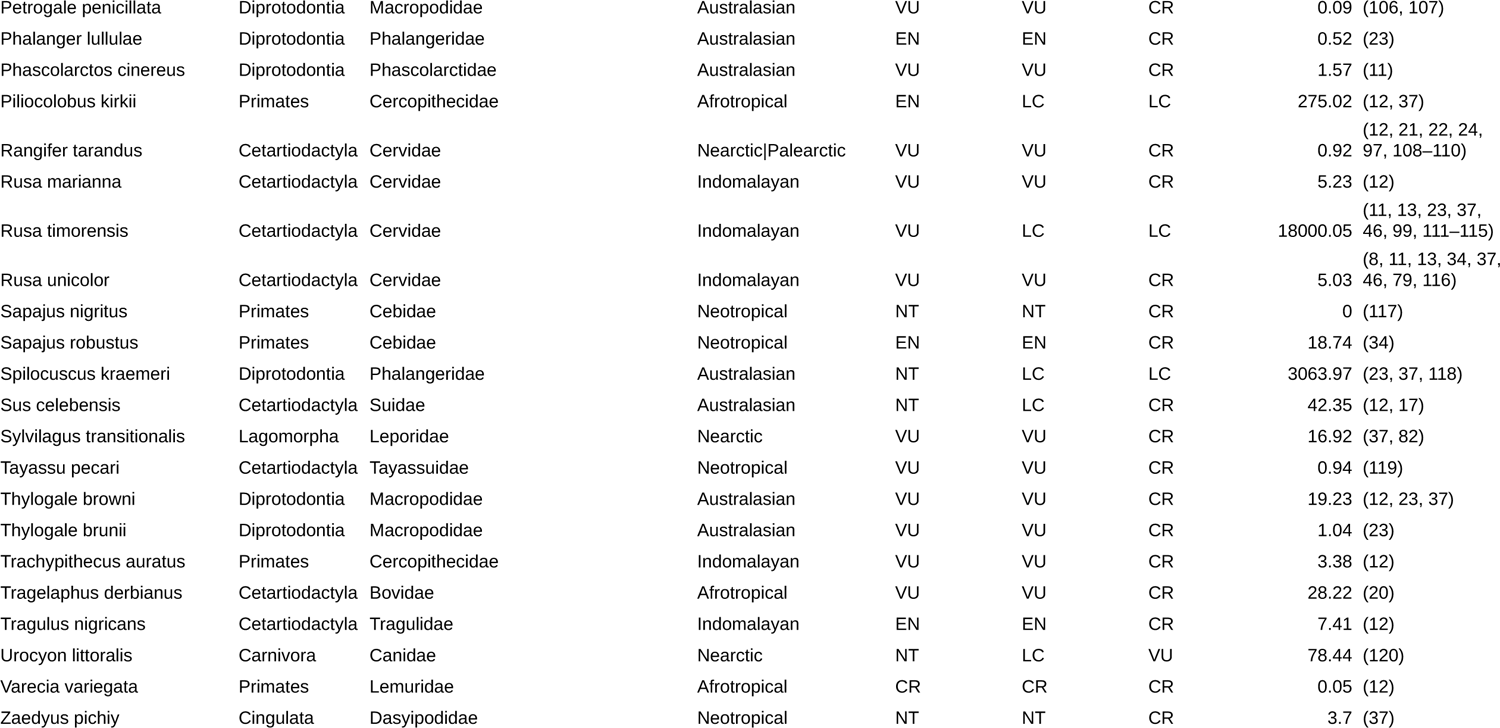
Migrant threatened mammal species included in analysis. Columns indicate species name, mammal family, whether the species is feral or not (indicated by feral nomenclature), their native realm (with multiple realms separated by ‘|’), their native-only threat status (i.e., current IUCN status), and threat statuses for the ‘expansive’ and ‘independent’ scenarios. ‘% Range Increase’ indicates the percent their range has increased via migration, which can equal infinity for species that are extinct in their native ranges.

**Fig. S1.**
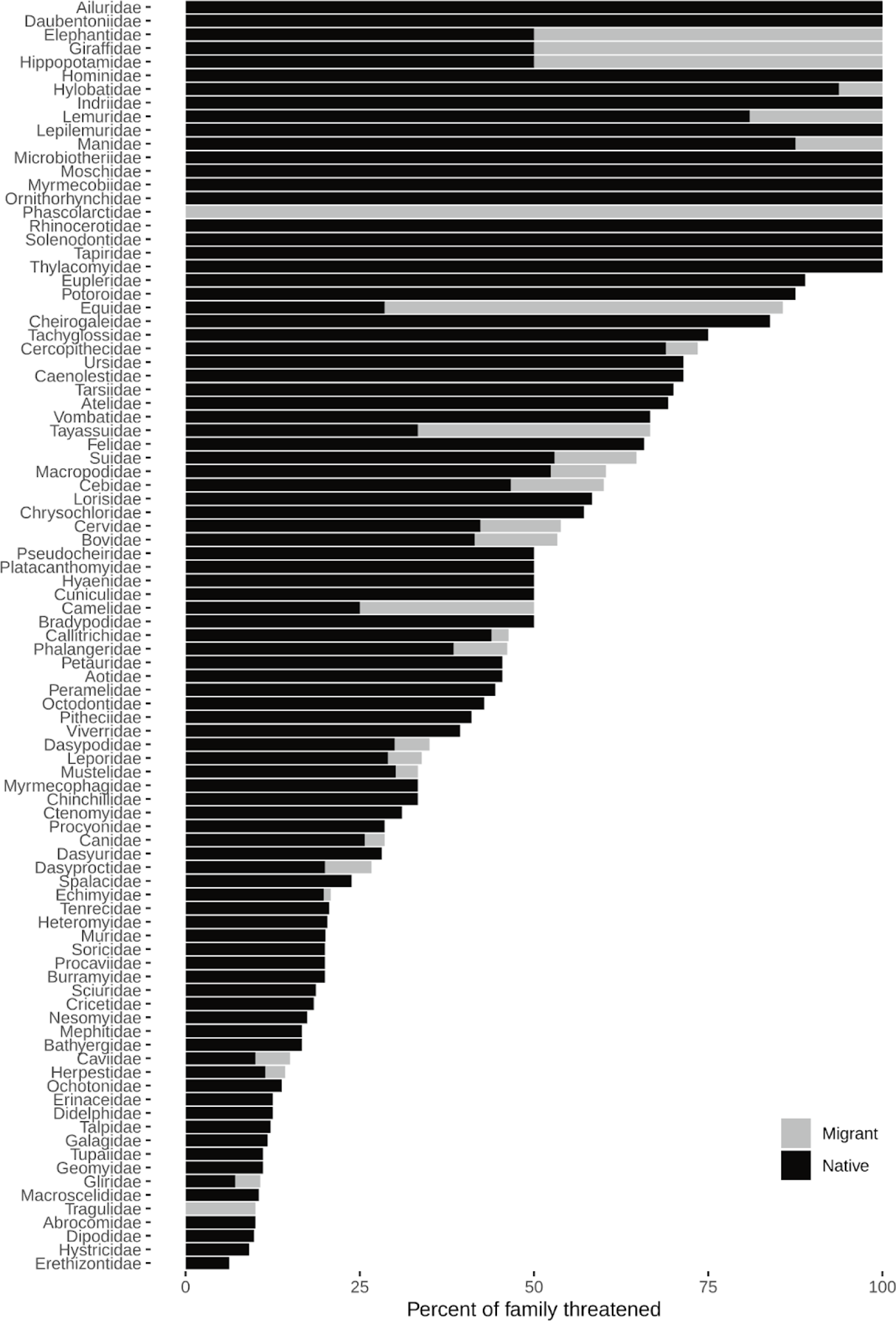
Percent of mammal species per family that are threatened and have migrant populations.

